# Combined deletion of cytosolic 5’-nucleotidases IA and II lowers glycemia by improving skeletal muscle insulin action and by lowering hepatic glucose production

**DOI:** 10.1101/2025.01.09.632106

**Authors:** R. Jacobs, G. Herinckx, N. Galland, C. Balty, D. Vertommen, M. H. Rider, M. Johanns

## Abstract

Obesity and type 2 diabetes (T2D)-linked hyperglycemia, along with their associated complications, have reached pandemic proportions, becoming a major public health issue. Genetic deletion or pharmacological inhibition of purine nucleotide-metabolizing enzymes has emerged as a potential strategy for treating diseases. We previously showed that cytosolic 5’-nucleotidase II (NT5C2)-deficient mice were protected against high-fat diet (HFD)-induced insulin resistance. In the present study, we investigated effects of dual deletion of cytosolic 5’-nucleotidase IA (NT5C1A) and NT5C2 in mice. We found that NT5C1A/NT5C2 double-knockout (NT5C-dKO) mice exhibited a hypoglycemic phenotype, displaying enhanced skeletal muscle insulin action and reduced hepatic glucose production. In addition to potential involvement of adenosine monophosphate (AMP)-activated protein kinase (AMPK) in their phenotype, NT5C-dKO mice displayed liver and skeletal muscle proteomic alterations most significantly linked to amino acid metabolism. Our findings support the development of novel anti-diabetic treatments using small-molecule cytosolic 5’-nucleotidase inhibitors.

## INTRODUCTION

Glycemia is determined by a balance between dietary intake, tissue uptake and utilization, and hepatic gluconeogenesis. After a meal, blood glucose levels rise, triggering insulin release from the pancreas. Insulin stimulates glucose uptake, primarily by skeletal muscle, while inhibiting hepatic glucose production. The net effect is a rapid return of glycemia to normal levels. However, type 2 diabetes (T2D) is characterized by insulin resistance due to a general loss of insulin sensitivity in cells and tissues. As a result, insulin in T2D fails to restore normal blood glucose levels after meals and to suppress hepatic glucose production, leading to hyperglycemia in both the postprandial and the starved states. AMP-activated protein kinase (AMPK) activation could be beneficial for treating diabetes by stimulating skeletal muscle glucose uptake [1,2] and perhaps by inhibiting hepatic gluconeogenesis, although the latter is somewhat controversial [3].

We and others have attempted targeting enzymes that metabolize purine nucleotides, specifically AMP and inosine 5’-monophosphate (IMP) as a means of achieving AMPK activation [4-9]. In most tissues, AMP is either directly hydrolyzed to adenosine by cytosolic 5’-nucleotidase IA (NT5C1A) or converted to IMP by AMP deaminase (AMPD). IMP can then be hydrolyzed to inosine by cytosolic 5’-nucleotidase II (NT5C2) (Figure 1A). NT5C1A expression is high in skeletal muscle but is low in liver, whereas NT5C2 is widely expressed, including in liver (Figure 1B, C).

**Figure 1:**
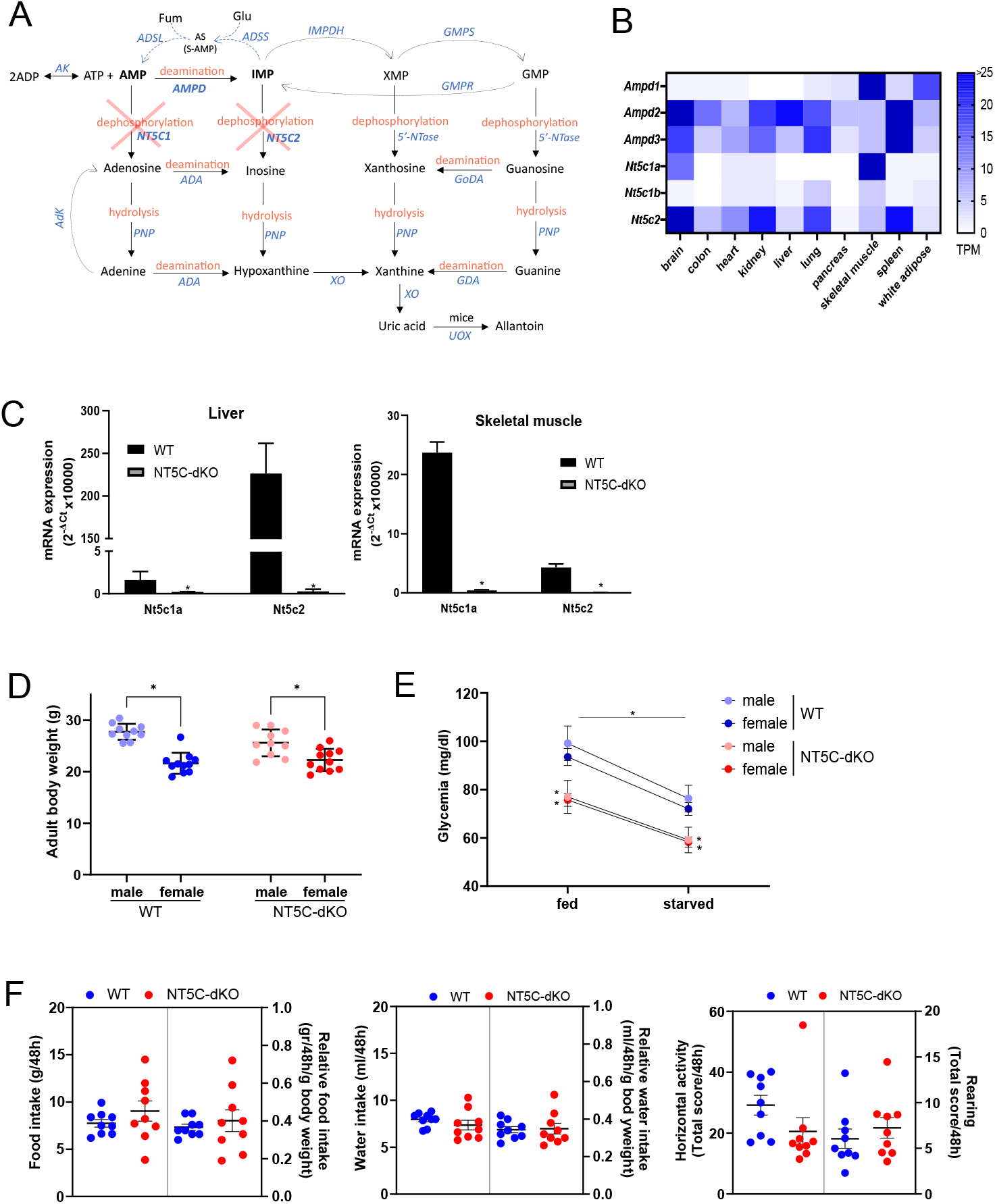
NT5C-dKO mice are mildly hypoglycemic. Purine nucleotide metabolism involves conversion of AMP to IMP by AMPDs and dephosphorylation of AMP to adenosine by NT5C1 or IMP to inosine by NT5C2 (**A**). IMP can be converted back to AMP via AS or directed towards the XMP/GMP pathways. AMP/IMP-metabolizing enzymes have distinct tissue distribution patterns. Data are for mouse tissues and were retrieved from the European Bioinformatics Institute (EBI) Expression Atlas (**B**). NT5C1A and NT5C2 mRNA levels were measured by RT-qPCR in skeletal muscle and liver from wild-type (WT) and NT5C1A-NT5C2 double-deficient (NT5C-dKO) mice to confirm deletion of the two enzymes (**C**). Adult male and female WT and NT5C-dKO mice were weighed (**D**). Glycemia was measured on blood drops from the tail of fed and overnight starved male and female WT and NT5C-dKO mice (**E**). WT and NT5C-dKO mice were monitored during 48 hours in physiocages for measurements of food consumption (left panel), water consumption (middle panel) and locomotor activity (right panel) (**F**).

In mice deficient AMPD1, the major isoenzyme in skeletal muscle, *ex vivo* muscle contraction induced AMP was seen, although this did not lead to increased glucose uptake [4]. Also, genetic deletion or pharmacological inhibition of AMPD1 failed to improve glycemic control in insulin-resistant HFD-fed mice fed [9]. Conversely deletion of the major liver isoenzyme AMPD2 had beneficial effects in mice fed a high-fructose diet, primarily by reducing hepatic glucose production [8].

Single whole-body knockout of either NT5C1A or NT5C2 did not lead to increased AMPK activation or contraction-induced glucose uptake in incubated skeletal muscles from mice chow-fed mice [6]. Our studies were at variance with those of others, who showed that invalidating NT5C1A or NT5C2 led to modestly increased skeletal muscle AMPKα Thr172 phosphorylation and glucose uptake [10]. Also, NT5C2 silencing was shown to activate AMPK in lung tumor cells [11]. However, we found that NT5C2-deficient mice were protected from HFD-induced weight gain, adiposity, insulin resistance, and hyperglycemia [7], with a possible implication of AMPK. Following up on these studies, here we created the first model of dual deletion of NT5C1A and NT5C2 (referred to as “NT5C-dKO” mice) and investigated effects on glycemic control, hepatic glucose production, insulin action in skeletal muscle and muscle/liver protein expression.

## METHODS

### Materials

Unless otherwise stated, all chemicals were from Merck/Sigma Aldrich. Radiochemicals were from Perkin Elmer. Antibodies were from the sources cited. Oligonucleotides were from Integrated DNA Technologies (Belgium). Compound “CRCD2” was from Enamine Ltd. (Ukraine, cat. no. 27358589).

### Animals

A whole-body NT5C1A:NT5C2 double-knockout C57Bl/6N mouse strain was obtained by crossing animals with single whole-body deletions of the two enzymes originally generated by AstraZeneca (Mölndal, Sweden) as previously described [6]. Homozygous animals were bred separately from the wild-type (WT) strain for up to three generations. NT5C-dKO mice exhibited no obvious phenotypic differences compared with WT mice and breeding was normal. Mice were housed with a 12 h light-dark cycle and fed *ad libitum* with a standard chow diet and water. Two-to eight-month-old males and females were used for experimentation and compared to age-matched WT animals. No obvious sex-based differences were observed comparing NT5C-dKO and WT mice. At the beginning of *in vivo* experiments, animals were weighed for normalization purposes. All experiments were approved by the Animal Ethics Committee of the UCLouvain under reference number 2021/UCL/MD/028 and conducted in accordance with EU Directive 2010/63/EU for animal experimentation.

### Bodyweight, food/water intake and locomotor activity

After a one-week adaptation period to re-housing, animals were kept for 72 hours in metabolic cages (Bioseb physiocage 00) for measurement of food/water consumption and horizontal movement over the last 48 hours at the UCLouvain Animal Behaviour Analysis Platform (BeAP). Animals were weighed at the start of the experiment for normalization.

### RNA extraction and RT-qPCR

Total RNA was extracted, mRNA was retrotranscribed to cDNA using poly-dT primers, and tissue expression of Nt5c1a and Nt5c2 was analyzed as previously described [7] using the following primers (Nt5c1a: forward CTCAGGTGGGAGTTCGTCTCA, reverse GGTAGCAGATGGGGCTATTCC; Nt5c2: forward TGACCGCTTACAGAATGCAG, reverse CGGCTAGGGTATAATCCATATCA) along with Rpl19 (forward GAAGGTCAAAGGGAATGTGTTCA, reverse CCTTGTCTGCCTTCAGCTTGT) as reference.

### Glucose, pyruvate and insulin tolerance tests

Overnight starved animals were subjected to an oral glucose tolerance test (OGTT) by gavage of 2 mg glucose/g of bodyweight, an intraperitoneal pyruvate tolerance test (ipPTT) by injection of 1 mg sodium pyruvate/g of bodyweight, or an insulin tolerance test by intraperitoneal injection of 0.1U of recombinant human insulin (Actrapid, Novo Nordisk). Glycemia was measured on blood drops taken from the tip of the tail at the indicated time intervals over 2 h using a glucometer (Free Style Lite, Abbott). The incremental area under the curve (iAUC) was calculated using the starting baseline of each group.

### Collection of organs and plasma

*Ad libitum* fed and overnight starved mice were euthanized to rapidly collect organs for subsequent analyses. Around 0.5 ml of blood was drawn by cardiac puncture and centrifuged (2000*g* x 10 min at 4

°C) to obtain plasma. The left lobe of the liver was immediately freeze-clamped to avoid ischemia. Skeletal muscles (gastrocnemius and/or soleus) were dissected for *ex vivo* incubation or frozen immediately for subsequent analyses.

### Incubation of skeletal muscles for measurements of glucose uptake and purine nucleotides or immunoblotting

Isolated soleus and gastrocnemius muscles from WT and NT5-dKO mice were incubated in parallel for 30 min with or without electrical stimulation or incubation with 100 nM insulin, for measurements of [^3^H]-2-deoxyglucose uptake or purine nucleotides as described [7]. Muscles were also incubated for immunoblotting with the indicated antibodies.

### Incubation of primary hepatocytes for measurements of glucose production and purine nucleotides or immunoblotting

Primary hepatocytes from WT and NT5C-dKO mice were isolated in parallel as described [12]. After plate attachment and overnight incubation, cells were either incubated for 2 h with 10 mM lactate plus 1 mM pyruvate in glucose-free media for measurements of glucose production, or in normal media containing glucose, for immunoblotting and measurement of purine nucleotides and metabolites by GC-MS as described [12,13]. Where indicated, 10 μM N-(3-carbamoyl-4,5,6,7-tetrahydrobenzo-[b]thiophen-2-yl)-1H-benzo[d]imidazole-5-carboxamide (“CRCD2”) [14] and 0.5 mM 5-Ethynyl-2-deoxyuridine (EdU) were included in the incubations to inhibit NT5C2 and NT5C1A, respectively.

### Pancreatic insulin release and content

Before (30 min) and after (30 min) administration of an oral bolus of glucose, blood was harvested for subsequent measurement of plasma insulin by ELISA (Crystal Chem, #90080) following the manufacturer’s instructions. For pancreatic insulin content, pancreata were collected from starved mice and processed as described [15] for ELISA measurement of insulin using the same kit.

### Liver and skeletal muscle proteomes

Whole tibialis anterior (TA) muscles or minced liver (∼100 mg ) were homogenized in 0.3 ml of lysis buffer containing 50 mM triethylammonium bicarbonate pH 8.5, 0.5% (w/v) sodium deoxycholate, 1% (v/v) Igepal, 0.1% (w/v) SDS, 0.2% (w/v) dodecyl maltoside, 1 mM dithiothreitol, 1 mM EDTA, 0.5 mM phenylmethylsulfonyl fluoride. Extracts were centrifuged (10 min x 20,000g at 4°C) and protein content was estimated by BCA assay (Thermo Scientific). Samples of muscle protein (100 μg) or liver protein (200 μg) were precipitated with 4 volumes of acetone overnight at -20°C. Pellets were washed and resuspended in 100 mM bicarbonate buffer (pH 8) for digestion with trypsin. Peptides were quantified (Pierce Quantitative Colorimetric Peptide Assay) and 10 μg of peptides per sample were taken for 16-plex tandem mass tag (TMT) labelling (Thermo Scientific) and pools were separated by a high pH reversed-phase peptide fractionation kit (Thermo Scientific) following the manufacturer’s instructions for in-house analysis by LC-MS/MS (Orbitrap Fusion Lumos MS, Thermo Scientific) as described [16]. Differential abundance was determined using the Proteome Discoverer 3.1 software (Thermo Scientific) and *P*_adj_<0.1 was judged significant. Enrichment analysis for associated gene ontology terms of biologic processes (GO-BP) and Kyoto Encyclopedia of Genes and Genomes (KEGG) pathways was carried out using Metascape (https://metascape.org/) with default parameters and GO-BP clusters were manually annotated.

### Statistical analyses

Unless otherwise specified, data were obtained from at least three separate experiments or from at least five animals per groups for comparison. Statistical analyses and graph construction were performed using Graphpad Prism 10 software. Statistical tests that were used are described in the figure legends and *P*<0.05 was judged significant.

## RESULTS

### Mice lacking both NT5C2 and NT5C1A have a mild hypoglycemic phenotype

Due to the lack of adequate antibodies for detecting NT5C1A and NT5C2 in tissue extracts, NT5C2 and NT5C1A expression was assessed at the mRNA level by qPCR. Expression of both Nt5c2 and Nt5c1a mRNA was quasi-totally absent in skeletal muscle and liver from NT5C-dKO versus WT mice (Figure 1C), indicating efficient deletion of both nucleotidases. There was no apparent phenotypic difference of male or female NT5C-dKO compared to WT mice, as body weight, food/water intake and locomotor activity were similar (Figure 1D,F). However, plasma glycemia was significantly lower in NT5C-dKO mice both in the fed state and after overnight starvation (Figure 1E), suggesting alterations in glycemic control.

### NT5C-dKO mice have improved glucose clearance through enhanced skeletal muscle insulin action

When subjected to an oral glucose tolerance test, the blood glucose concentrations were significantly lower in NT5C-dKO compared to WT mice and the area under the curve (AUC) over 2h following glucose gavage was significantly reduced in NT5C-dKO mice (Figure 2A).

**Figure 2:**
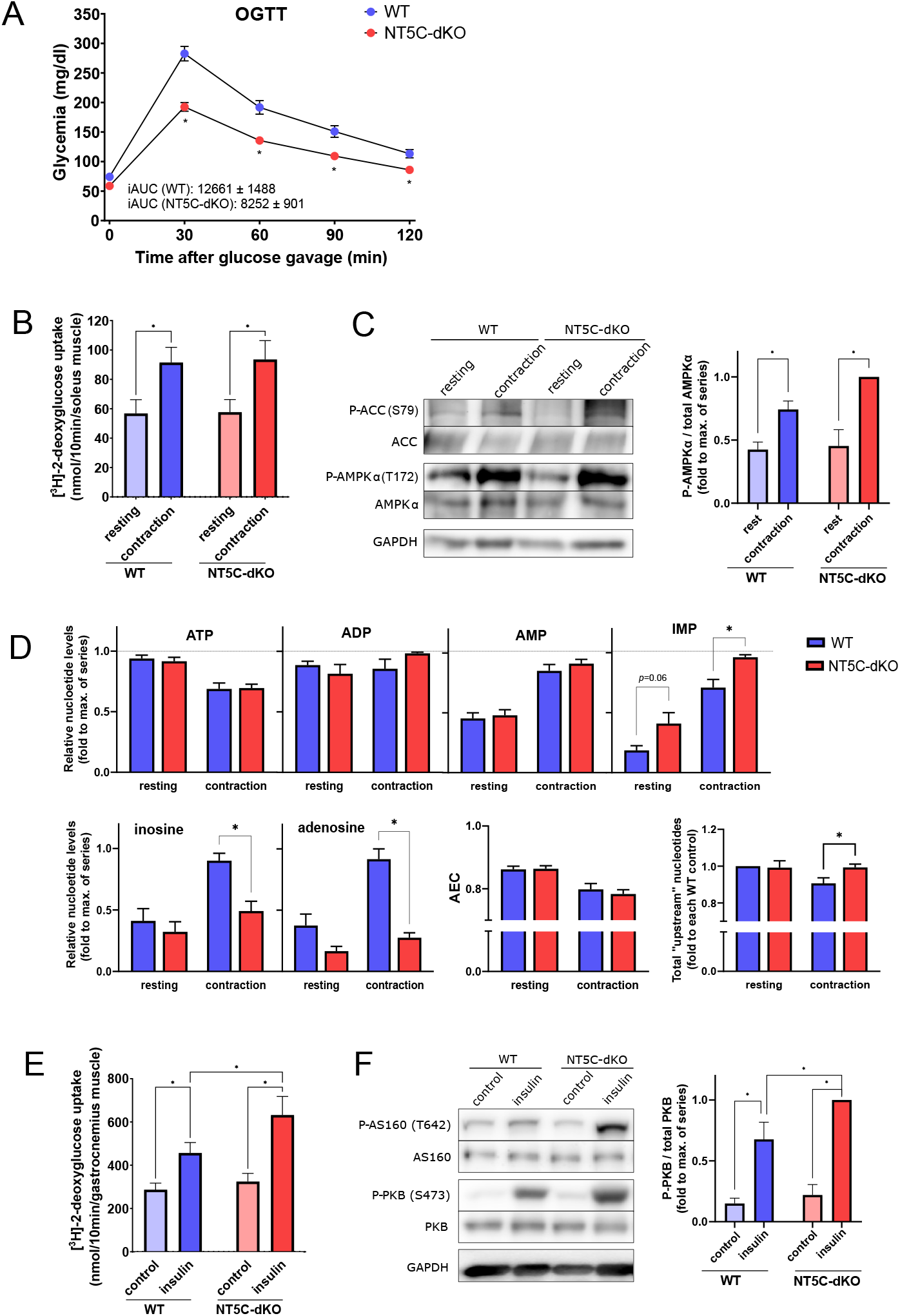
Insulin-stimulated glucose uptake is increased in muscles from NT5C-dKO mice. Starved WT and NT5C-dKO mice were subjected to an oral glucose tolerance test (OGTT, 2 mg glucose/g of body weight) and glycemia was measured on blood drops from the tail at the indicated times over 2h (**A**). Isolated soleus muscles from WT and NT5C-dKO mice were incubated *ex vivo* for 30 min, either with or without electrically stimulated contraction (**B-D**) or in the presence or absence of 100 nM insulin (**E, F**) for measurement of [^3^H]-2-deoxyglucose uptake during an additional 10 min of incubation (**B, E**), immunoblotting with the indicated antibodies (**C, F**) or measurement of purine nucleotides in perchloric acid extracts by HPLC (**D**). In D, “upstream” nucleotides are ATP+ADP+AMP+IMP.

As glucose removal primarily occurs via uptake into skeletal muscle, which is stimulated both by contraction and insulin, we looked whether this process was altered in NT5C-dKO mice. *Ex vivo* contraction of soleus muscles resulted in increased [^3^H]-2-deoxyglucose uptake as expected (Figure 2B) but was similar in muscles from WT and NT5C-dKO mice. Measurements of intramuscular purine nucleotide concentrations indicated that contraction-induced increases in AMP were similar in soleus muscles from WT versus NT5C-dKO mice, whereas the rise in IMP was potentiated (Figure 2D). Immunoblotting of muscle extracts indicated similar contraction-induced increases in phosphorylation of AMPKα Thr172 and ACC Ser212 (Figure 2C), a direct AMPK target, suggesting there was no difference in AMPK activation due to contraction in skeletal muscle from WT versus NT5C-dKO mice. By contrast, insulin-stimulated [^3^H]-2-deoxyglucose uptake was significantly enhanced in *ex vivo* incubations of gastrocnemius muscles from NT5C-dKO versus WT mice (Figure 2E). Accordingly, insulin-induced PKB phosphorylation was potentiated in extracts of muscles from NT5C-dKO mice compared to WT mice (Figure 2F), suggesting improved muscle insulin sensitivity due to dual deletion of NT5C1A and NT5C2. Therefore, an intraperitoneal insulin tolerance test on WT versus NT5C-dKO mice was undertaken. Using a submaximal dose of insulin, glycemia was significantly reduced in NT5C-dKO versus WT mice (Supplementary Figure S1A). In addition to improved systemic insulin responsiveness, pancreatic insulin release following glucose gavage appeared to be enhanced in NT5C-dKO compared with WT mice (Supplementary Figure S1B), which could not be explained by increased pancreatic insulin content (Supplementary Figure S1C), suggesting improved pancreatic insulin release in response to glucose in NT5C-dKO mice.

### Dual NT5C1A/NT5C2 deletion lowers hepatic glucose production

As hepatic glucose production is the main determinant of glycemia during starvation, we looked whether the observed differences in blood sugar after starvation could be due to reduced glucose synthesis. This was indeed suggested by an intraperitoneal pyruvate tolerance test (ipPTT), during which the rise in glycemia was significantly lower in NT5C-dKO compared to WT mice (Figure 3A). Accordingly, the metabolic stress-induced reduction in glucose production was more pronounced in hepatocytes from NT5C-dKO compared to WT mice (Figure 3B). In these experiments, metabolic poisons were used to increase the flux through the purine nucleotide system, namely phenformin, which inhibits mitochondrial complex I or oligomycin, which inhibits mitochondrial ATP synthase. The net result is a decrease in mitochondrial ATP production causing a concomitant rise in AMP. AMPKα Thr172 phosphorylation was slightly higher in NT5C-dKO versus WT hepatocytes incubated with both compounds (Figure 3C). In fact, physiological energy stress is encountered in liver during long-term starvation [17], accompanied by a drop in ATP and a rise in AMP levels. In hepatocytes, AMP and IMP are potent negative regulators of glucose production, by inhibiting fructose-1,6-bisphophatase (FBPase-1) catalyzing a key step of gluconeogenesis [18]. Interestingly, levels of IMP but not of AMP were significantly enhanced in hepatocytes from NT5C-dKO versus WT mice incubated with oligomycin or phenformin (Figure 3D), which might have been due to rapid conversion of AMP to IMP by AMPD. Under these conditions, nucleotides can be rerouted towards the TCA cycle through a salvage pathway involving conversion of IMP plus glutamate to adenylosuccinate (AS) and fumarate by AS synthase (ADSS) and back to AMP by AS lyase (ADSL) (Figure 1A). Indeed by GC-MS, glutamate levels were decreased in NT5C-dKO versus WT hepatocytes, while fumarate and malate were increased in presence of phenformin (Supplementary Figure S2).

**Figure 3:**
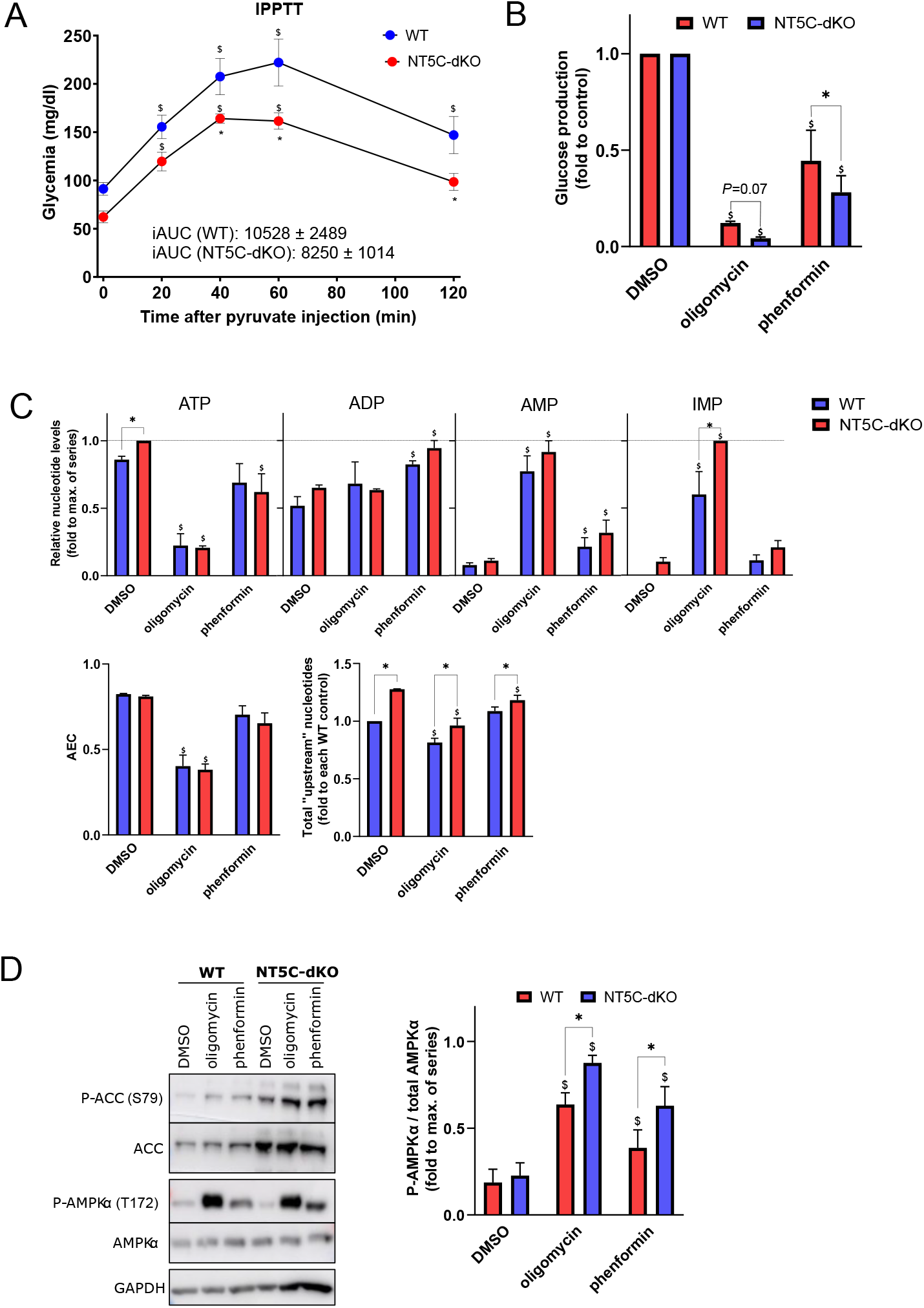
NT5C-dKO mice display decreased hepatic glucose production. Starved WT and NT5C-dKO mice were subjected to an intraperitoneal pyruvate tolerance test (IPPTT, 1 mg pyruvate/g of body weight) and glycemia was measured on blood drops from the tail at the indicated intervals over two hours (**A**). Hepatocytes were isolated from the livers of WT and NT5C-dKO mice, cultured overnight and incubated for 2 h with or without 1 uM oligomycin or 0.5 mM phenformin, either in glucose-free media supplemented with 10 mM lactate and 1 mM pyruvate for enzymatic determination of newly produced glucose in the media (**B**), or in normal media containing glucose for immunoblotting with the indicated antibodies (**C**) or measurement of purine nucleotides in perchloric acid cell extracts by HPLC (**D**). In D, “upstream” nucleotides are ATP+ADP+AMP+IMP.

### Muscle and liver proteomes are remodeled in NT5C-dKO mice

To investigate possible mechanisms underlying the changes in the regulation of skeletal muscle glucose uptake and liver glucose production, we analyzed the proteome in both tissues by LC-MS/MS. Using TMT labelling, we were able to robustly quantify over 3,000 proteins (Figure 4A,B). Differences in abundance of 49 and 181 proteins (*P*_adj_.<0.1) were seen in NT5C-dKO versus WT muscles and livers, respectively (Figure 4A,B & Supplementary Table S1). Interestingly, AS synthase was among the proteins slightly more abundant in NT5C-dKO versus WT livers. Functional analysis of associated GO-BP and KEGG terms revealed that these differential proteins were enriched for nucleotide metabolism, as expected, but also strongly for amino acid metabolism. Indeed, (purine) nucleotide and amino acid metabolism have multiple connections at the level of *de novo* synthesis, catabolism and salvage pathways [19]. Together, these data indicate that NT5C-dKO mice undergo a subtle proteomic remodeling, possibly to reroute nucleotide catabolism.

**Figure 4:**
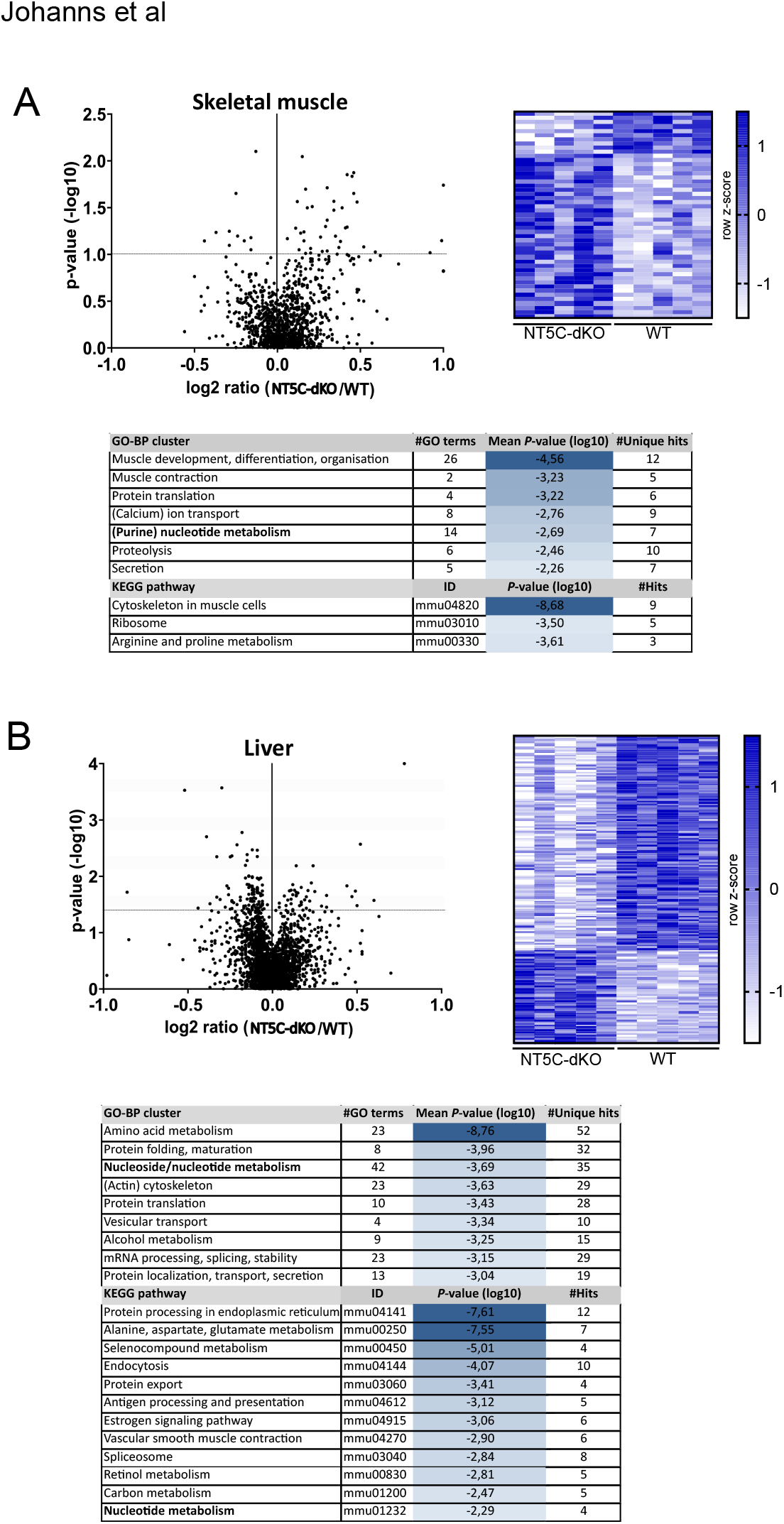
Proteomic remodeling in muscle and liver of NT5C-dKO mice. Tibialis anterior (TA) muscles (**A**) and livers (**B**) from WT and NT5C-dKO mice were collected for protein extraction, trypsin digestion, TMT peptide labelling and proteomic analysis by LC-MS/MS. Differentially abundant proteins (*P*_adj_.<0.1) are highlighted in volcano plots and represented in heatmaps for each tissue, with functional enrichment analysis for GO-BP and KEGG pathway terms. GO-BP terms are manually annotated showing clusters containing the indicated number of individual GO terms along with their mean *P*-values calculated using the Metascape online tool.

### Pharmacological NT5C2/NT5C1 inhibition decreases hepatocyte glucose production

To explore the therapeutic potential of our findings, we attempted to pharmacologically mimic dual deletion of NT5C1A and NT5C2, by the combined use of inhibitors on cells/tissues from WT mice. The nucleotide analog 5-ethynyl-2′-deoxyuridine (EdU) has been used to inhibit NT5C1A [20], while the recently developed small-molecule inhibitor “CRCD2” inhibits NT5C2 [14]. In *ex vivo* incubated skeletal muscles, both basal and contraction-induced glucose uptake and nucleotide levels were unaltered by incubation with both inhibitors (data not shown). We therefore tested their effect with longer incubation times in primary hepatocytes, a more flexible model. When metabolic stress was induced by oligomycin or phenformin treatment, AMP and IMP levels increased to higher levels in the presence of NT5C inhibitors (Figure 5A). Accordingly, the phenformin and oligomycin-induced inhibition of glucose production was stronger in presence of NT5C inhibitors (Figure 5B).

**Figure 5:**
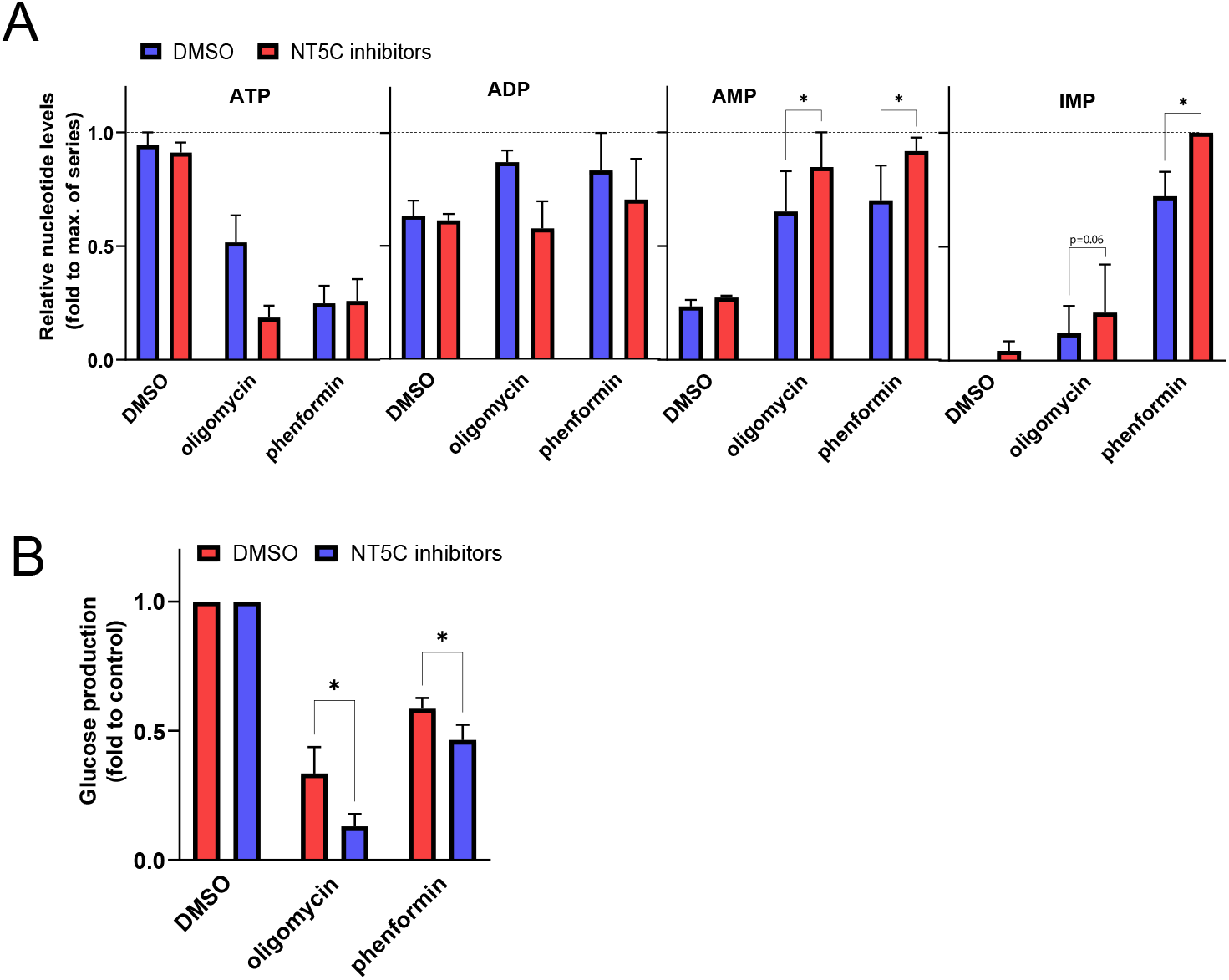
Pharmacological inhibition of NT5C1 and NT5C2 in hepatocytes recapitulates genetic deletion. Hepatocytes were isolated from the livers of WT and NT5C-dKO mice, cultured overnight and incubated for 4 h in glucose-free media supplemented with 10 mM lactate and 1 mM pyruvate with or without 0.5 uM oligomycin or 0.5 mM phenformin, alone or in combination with 0.5 mM EdU and 10 uM CRCD2 to inhibit NT5C1 and NT5C2, respectively, for measurement of purine nucleotides in perchloric acid cell extracts by HPLC (**A**) or enzymatic determination of glucose in incubation media (**B**).

## DISCUSSION

T2D-linked hyperglycemia is a major global public health issue and predisposes affected individuals to metabolic and cardiovascular disease. Existing treatments have the downside of potentially causing the development of resistance and/or intolerance. Increased cellular purine dephosphorylation is a feature seen in both type 1 and type 2 diabetic patients [21], while plasma hypoxanthine [22], uric acid, xanthine and adenosine [23] were positively correlated to the incidence of T2D. In the present study, we propose an alternative strategy to lower glycemia, by dual inhibition of soluble NT5Cs. Overall, we found that purine nucleotide metabolism was remarkably resilient towards metabolic stress or genetic manipulation, probably because fluxes through the enzymes are very low and because, from IMP and glutamate, adenine nucleotides could be rerouted towards the TCA cycle via ADSS/ADSL-mediated fumarate production (Figure 1A), with other possible connections to amino acid metabolism through the TCA cycle . Nevertheless, NT5C-dKO mice displayed mild fasting hypoglycemia and improved glucose clearance by mechanisms of i) improved skeletal muscle insulin action, ii) enhanced stress-induced suppression of hepatic glucose production, and iii) higher glucose-induced pancreatic insulin release. We cannot rule out a role for AMPK activation in the hypoglycemic phenotype seen upon dual NT5C1A-NT5C2 deletion. AMPK activation *in vivo* is difficult to study because once tissues go anoxic during dissection prior to freeze-clamping, AMPK becomes activated. Of interest in explaining effects seen in NT5C-dKO mice is the enhanced rise in IMP seen during muscle contraction (Figure 2D) and in hepatocytes incubated with mitochondrial poisons (Figure 5A). We estimate that the maximal concentrations that would be reached in skeletal muscle and hepatocytes would be around 500 nmol/g of wet weight and 5 nmol/10^6^ cells, respectively. Although IMP does not appear to, at least directly, allosterically stimulate AMPK activity nor protect against AMPKα Thr172 dephosphorylation [24], IMP is an allosteric inhibitor of FBPase-1 [18], which might participate in enhanced suppression of hepatic glucose production due to oligomycin or phenformin treatment (Figure 5B). Another enzyme that is IMP-sensitive is the phosphomannomutase isoenzyme (PMM1) that hydrolyses glucose-1,6-bisphosphate [25], raising the intriguing possibility that reduced glucose-1,6-bisphosphate levels might be linked to the control of glycemia. It is perhaps noteworthy that inhibition of IMP dehydrogenase reduces diet-induced obesity [26]. Lastly, dual NT5C inhibition might increase flux through the XMP pathway (Figure 1A) and hypoxanthine supplementation was recently shown to decrease glycemia by suppressing hepatic gluconeogenesis [27].

Our findings support the development of new anti-diabetic compounds based on small-molecule inhibition of cytosolic NT5Cs. In fact, NT5C inhibitors are already being tested in anti-cancer therapy [14] and might be repurposed to treat T2D. Despite the absence of any apparent adverse health effects of whole-body deletion of both NT5C1A and NTC2 in our mouse model, the presence of auto-antibodies against NT5C1A were linked to (dermato)myositis [28] and loss-of-function mutations of NT5C2 were associated with spastic paraplegia [29], while gain-of-function mutations in NT5C2 were linked to persistence of acute lymphoid leukemia (ALL) [30]. Long-term, partial inhibition of NT5Cs is likely to be beneficial without major side-effects. In any event, there is an increasing need for the development of novel, isoform-specific small-molecule inhibitors of NT5Cs. One option to achieve this could be to target functionally essential enzyme oligomerization, a strategy that was successfully employed by our colleagues to disrupt LDH tetramers required by cancer cells [31], and both NT5C1A and NT5C2 are active as homotetramers [32].

## REFERENCES

1. Esquejo, R.M., Albuquerque, B., Sher, A., Blatnik, M., Wald, K., Peloquin, M., et al. (2022) AMPK activation is sufficient to increase skeletal muscle glucose uptake and glycogen synthesis but is not required for contraction-mediated increases in glucose metabolism. Heliyon 8:e11091, doi:10.1016/j.heliyon.2022.e11091

2. Jensen, T.E., Wojtaszewski, J.F., Richter, E.A. (2009) AMP-activated protein kinase in contraction regulation of skeletal muscle metabolism: necessary and/or sufficient? Acta Physiol. (Oxf.) 196:155–174, doi:10.1111/j.1748-1716.2009.01979.x

3. Johanns, M., Hue, L., Rider, M.H. (2023) AMPK inhibits liver gluconeogenesis: fact or fiction? Biochem. J. 480:105–125, doi:10.1042/bcj20220582

4. Plaideau, C., Lai, Y.C., Kviklyte, S., Zanou, N., Lofgren, L., Andersen, H., et al. (2014) Effects of pharmacological AMP deaminase inhibition and Ampd1 deletion on nucleotide levels and AMPK activation in contracting skeletal muscle. Chem Biol 21:1497–1510, doi:10.1016/j.chembiol.2014.09.013

5. Plaideau, C., Liu, J., Hartleib-Geschwindner, J., Bastin-Coyette, L., Bontemps, F., Oscarsson, J., et al. (2012) Overexpression of AMP-metabolizing enzymes controls adenine nucleotide levels and AMPK activation in HEK293T cells. FASEB J. 26:2685–2694, doi:10.1096/fj.11-198168

6. Kviklyte, S., Vertommen, D., Yerna, X., Andersen, H., Xu, X., Gailly, P., et al. (2017) Effects of genetic deletion of soluble 5’-nucleotidases NT5C1A and NT5C2 on AMPK activation and nucleotide levels in contracting mouse skeletal muscles. Am. J. Physiol. Endocrinol. Metab. 313:E48–e62, doi:10.1152/ajpendo.00304.2016

7. Johanns, M., Kviklyte, S., Chuang, S.J., Corbeels, K., Jacobs, R., Herinckx, G., et al. (2019) Genetic deletion of soluble 5’-nucleotidase II reduces body weight gain and insulin resistance induced by a high-fat diet. Mol. Genet. Metab. 126:377–387, doi:10.1016/j.ymgme.2019.01.017

8. Hudoyo, A.W., Hirase, T., Tandelillin, A., Honda, M., Shirai, M., Cheng, J., et al. (2017) Role of AMPD2 in impaired glucose tolerance induced by high fructose diet. Mol. Genet. Metab. Rep. 13:23–29, doi:10.1016/j.ymgmr.2017.07.006

9. Admyre, T., Amrot-Fors, L., Andersson, M., Bauer, M., Bjursell, M., Drmota, T., et al. (2014) Inhibition of AMP deaminase activity does not improve glucose control in rodent models of insulin resistance or diabetes. Chem Biol 21:1486–1496, doi:10.1016/j.chembiol.2014.09.011

10. Kulkarni, S.S., Karlsson, H.K., Szekeres, F., Chibalin, A.V., Krook, A., Zierath, J.R. (2011) Suppression of 5’-nucleotidase enzymes promotes AMP-activated protein kinase (AMPK) phosphorylation and metabolism in human and mouse skeletal muscle. J. Biol. Chem. 286:34567–34574, doi:10.1074/jbc.M111.268292

11. Pesi, R., Allegrini, S., Garcia-Gil, M., Piazza, L., Moschini, R., Jordheim, L.P., et al. (2021) Cytosolic 5’-Nucleotidase II Silencing in Lung Tumor Cells Regulates Metabolism through Activation of the p53/AMPK Signaling Pathway. Int. J. Mol. Sci. 22, doi:10.3390/ijms22137004

12. Johanns, M., Corbet, C., Jacobs, R., Drappier, M., Bommer, G.T., Herinckx, G., et al. (2022) Inhibition of basal and glucagon-induced hepatic glucose production by 991 and other pharmacological AMPK activators. Biochem. J. 479:1317-1336, doi:10.1042/bcj20220170

13. Johanns, M., Lai, Y.C., Hsu, M.F., Jacobs, R., Vertommen, D., Van Sande, J., et al. (2016) AMPK antagonizes hepatic glucagon-stimulated cyclic AMP signalling via phosphorylation-induced activation of cyclic nucleotide phosphodiesterase 4B. Nat. Commun. 7:10856, doi:10.1038/ncomms10856

14. Reglero, C., Dieck, C.L., Zask, A., Forouhar, F., Laurent, A.P., Lin, W.W., et al. (2022) Pharmacologic Inhibition of NT5C2 Reverses Genetic and Nongenetic Drivers of 6-MP Resistance in Acute Lymphoblastic Leukemia. Cancer Discov. 12:2646–2665, doi:10.1158/2159-8290.Cd-22-0010

15. Clee, S.M., Yandell, B.S., Schueler, K.M., Rabaglia, M.E., Richards, O.C., Raines, S.M., et al. (2006) Positional cloning of Sorcs1, a type 2 diabetes quantitative trait locus. Nat. Genet. 38:688–693, doi:10.1038/ng1796

16. Baes, R., Grünberger, F., Pyr Dit Ruys, S., Couturier, M., De Keulenaer, S., Skevin, S., et al. (2023) Transcriptional and translational dynamics underlying heat shock response in the thermophilic crenarchaeon Sulfolobus acidocaldarius. mBio 14:e0359322, doi:10.1128/mbio.03593-22

17. Hasenour, C.M., Ridley, D.E., James, F.D., Hughey, C.C., Donahue, E.P., Viollet, B., et al. (2017) Liver AMP-Activated Protein Kinase Is Unnecessary for Gluconeogenesis but Protects Energy State during Nutrient Deprivation. PLoS One 12:e0170382, doi:10.1371/journal.pone.0170382

18. Hunter, R.W., Hughey, C.C., Lantier, L., Sundelin, E.I., Peggie, M., Zeqiraj, E., et al. (2018) Metformin reduces liver glucose production by inhibition of fructose-1-6-bisphosphatase. Nat. Med. 24:1395–1406, doi:10.1038/s41591-018-0159-7

19. Stenesh, J. Amino Acid and Nucleotide Metabolism. In: Stenesh J, editor. Biochemistry. Boston, MA: Springer US; 1998. p. 345–374.

20. Garvey, E.P., Prus, K.L. A specific inhibitor of heart cytosolic 5’-nucleotidase I attenuates hydrolysis of adenosine 5’-monophosphate in primary rat myocytes.

21. Dudzinska, W. (2014) Purine nucleotides and their metabolites in patients with type 1 and 2 diabetes mellitus. Journal of Biomedical Science and Engineering 7:38–44, doi:10.4236/jbise.2014.71006.

22. Ottosson, F., Smith, E., Gallo, W., Fernandez, C., Melander, O. (2019) Purine Metabolites and Carnitine Biosynthesis Intermediates Are Biomarkers for Incident Type 2 Diabetes. J Clin Endocrinol Metab 104:4921–4930, doi:10.1210/jc.2019-00822

23. Papandreou, C., Li, J., Liang, L., Bulló, M., Zheng, Y., Ruiz-Canela, M., et al. (2019) Metabolites related to purine catabolism and risk of type 2 diabetes incidence; modifying effects of the TCF7L2-rs7903146 polymorphism. Sci. Rep. 9:2892, doi:10.1038/s41598-019-39441-6

24. Toyoda, T., Hayashi, T., Miyamoto, L., Yonemitsu, S., Nakano, M., Tanaka, S., et al. (2004) Possible involvement of the alpha1 isoform of 5’AMP-activated protein kinase in oxidative stress-stimulated glucose transport in skeletal muscle. Am. J. Physiol. Endocrinol. Metab. 287:E166–173, doi:10.1152/ajpendo.00487.2003

25. Veiga-da-Cunha, M., Vleugels, W., Maliekal, P., Matthijs, G., Van Schaftingen, E. (2008) Mammalian phosphomannomutase PMM1 is the brain IMP-sensitive glucose-1,6-bisphosphatase. J. Biol. Chem. 283:33988–33993, doi:10.1074/jbc.M805224200

26. Su, H., Gunter, J.H., de Vries, M., Connor, T., Wanyonyi, S., Newell, F.S., et al. (2009) Inhibition of inosine monophosphate dehydrogenase reduces adipogenesis and diet-induced obesity. Biochem Biophys Res Commun 386:351–355, doi:10.1016/j.bbrc.2009.06.040

27. Huang, S., Liang, H., Chen, Y., Liu, C., Luo, P., Wang, H., Du, Q. (2024) Hypoxanthine ameliorates diet-induced insulin resistance by improving hepatic lipid metabolism and gluconeogenesis via AMPK/mTOR/PPARα pathway. Life Sci. 357:123096, doi:10.1016/j.lfs.2024.123096

28. Amlani, A., Choi, M.Y., Tarnopolsky, M., Brady, L., Clarke, A.E., Garcia-De La Torre, I., et al. (2019) Anti-NT5c1A Autoantibodies as Biomarkers in Inclusion Body Myositis. Front. Immunol. 10:745, doi:10.3389/fimmu.2019.00745

29. Darvish, H., Azcona, L.J., Tafakhori, A., Ahmadi, M., Ahmadifard, A., Paisán-Ruiz, C. Whole genome sequencing identifies a novel homozygous exon deletion in the NT5C2 gene in a family with intellectual disability and spastic paraplegia. LID - 20 [pii] LID - 10.1038/s41525-017-0022-7 [doi].

30. Dieck, C.L., Ferrando, A. (2019) Genetics and mechanisms of NT5C2-driven chemotherapy resistance in relapsed ALL. Blood 133:2263–2268, doi:10.1182/blood-2019-01-852392

31. Thabault, L., Brustenga, C., Savoyen, P., Van Gysel, M., Wouters, J., Sonveaux, P., et al. (2022) Discovery of small molecules interacting at lactate dehydrogenases tetrameric interface using a biophysical screening cascade. Eur. J. Med. Chem. 230:114102, doi:10.1016/j.ejmech.2022.114102

32. Walldén, K., Nordlund, P. (2011) Structural basis for the allosteric regulation and substrate recognition of human cytosolic 5’-nucleotidase II. J. Mol. Biol. 408:684–696, doi:10.1016/j.jmb.2011.02.059

