## Supplemental Figure S1 for "Combined deletion of cytosolic 5’-nucleotidases IA and II lowers glycemia by improving skeletal muscle insulin action and by lowering hepatic glucose production"

### Suppl. Figure S1

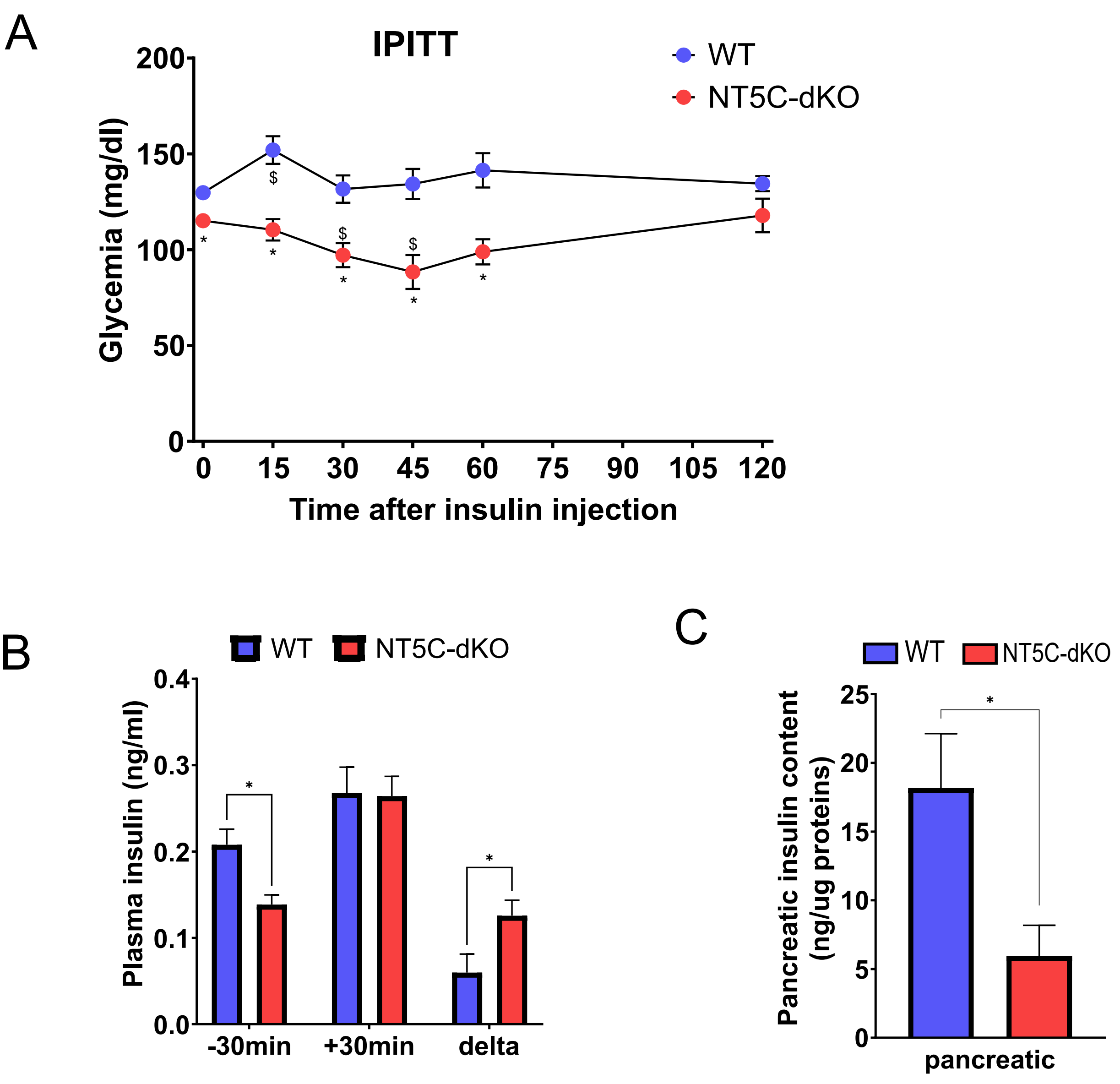


**Supplemental Figure S1: Enhanced insulin sensitivity, glucose-induced secretion but not pancreatic content in NT5C-dKO mice.** 4h-starved mice were subjected to an intraperitoneal insulin (0.1 U) tolerance test and glycemia was measured at indicated intervals (A). Blood was harvested before and after an oral bolus of glucose for measurement of plasma insulin by ELISA (B). Pancreatic insulin content of over-night starved mice was measured by ELISA in acid-ethanol extracts. Data are means ± s.e.m.from 5-8 animals per group and * indicates a significant (p<) difference (2-way ANOVA with Fisher's post-hoc test).
