## Supplemental Figure S2 for "Combined deletion of cytosolic 5’-nucleotidases IA and II lowers glycemia by improving skeletal muscle insulin action and by lowering hepatic glucose production"

### Suppl. Figure S2

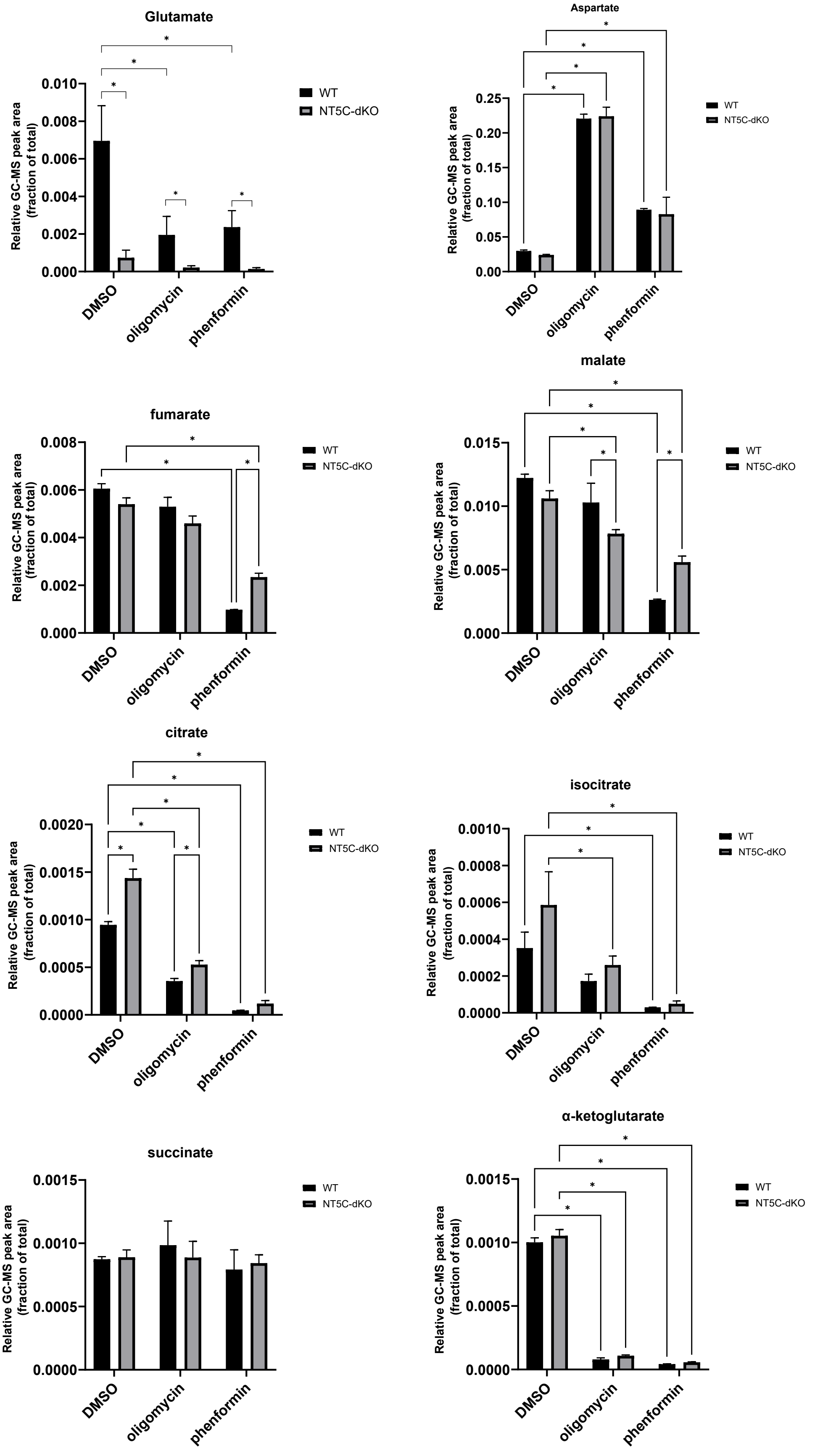


**Supplemental Figure S2: Altered TCA cycle and amino acid usage in NT5C-dKO mice.** Over-night cultured hepatocytes from WT versus NT5C-dKO mice were incubated for 2 hours in normal media supplemented with DMSO (vehicle control), 1 uM oligomycin or 0.5 mM phenformin. Polar metabolites were extracted using methanol-chloroform and derivatized by methoxyamine and MSTFA for analysis by GC-MS. Data are means ± s.e.m. of 5 biological replicates and * indicates a significant ($p<0.05$) difference (2-way ANOVA with Fisher's post-hoc test).
